# Resampling-based validation of a SNP panel for hybrid detection across generations: a case study in European lobster

**DOI:** 10.64898/2026.01.20.700654

**Authors:** Erik Sandertun Røed, Charlie Ellis, Jamie R. Stevens, Louise Chavarie, Marie Saitou

**Affiliations:** Centre for Ecological and Evolutionary Synthesis (CEES), University of Oslo (UiO), Blindernveien 31, Entrance: Moltke Moes vei, NO-0371 Oslo, Norway; Faculty of Biosciences, Norwegian University of Life Sciences (NMBU), Oluf Thesens vei 6, 1433 Ås, Norway; Department of Biosciences, University of Exeter, Prince of Wales Road, Exeter EX2 4PS, United Kingdom; Faculty of Environmental Sciences and Natural Resource Management, Norwegian University of Life Sciences (NMBU), Høgskoleveien 12, 1433 Ås, Norway

**Keywords:** hybridization, introgression, SNP panel, resampling, conservation genetics, fishery management

## Abstract

Accurate detection of hybridization and introgression is critical for both evolutionary research and applied conservation. In many systems, however, hybrid ancestry is difficult to detect beyond the F1 generation, especially when based on limited genetic markers. In European waters, hybridization between the native *Homarus gammarus* and the invasive *H. americanus* poses a direct risk to the integrity of native stocks and effective fishery management, yet detection methods are often limited to morphological traits or first-generation hybrids. A set of 79 SNPs previously developed to distinguish species between American and European lobsters and F1 individuals has shown promise, but its capacity to resolve later-generation backcrosses remains untested. Here, we present a resampling-based evaluation of this panel’s performance under realistic introgression scenarios, using individual-based population genetic models informed by empirical data. We show that the panel retains discriminatory power across multiple hybrid classes, with diminishing accuracy in second-generation backcrosses. These findings validate the panel’s utility for applied monitoring and highlight the broader potential of resumpling-anchored frameworks to benchmark hybrid detection tools in a wide range of species.

**Article summary:** This study tests how well a reduced panel of genetic markers can detect hybridization across multiple generations. Using empirical genetic data of a 79-SNP panel from European and American lobsters, the authors generated individuals with known ancestry proportions through a resampling framework that preserves observed genetic variation. These data were analysed using model-based genetic assignment and ordination. The results show that the marker panel reliably identifies pure species and first-generation hybrids, but has reduced power to distinguish later backcross generations, mainly between adjacent hybrid classes. The study provides a practical benchmark for evaluating reduced marker panels used in applied monitoring and conservation genetics.

## Introduction

Accurate identification of species boundaries is essential for understanding gene flow, reproductive isolation, and the ecological consequences of admixture (Chan et al. 2017; Christie and Strauss 2019; Satokangas et al. 2023; Wu et al. 2024; Chambers et al. 2025). In many taxa, however, species boundaries are obscured by cryptic genetic divergence or by interspecific hybridization, particularly following secondary contact between closely related species (Sardell and Uy 2016; Porto-Hannes et al. 2021; Termignoni-Garcia et al. 2022; Peng et al. 2025).

Cryptic divergence, in which genetically distinct lineages show little or no morphological differentiation, complicates taxonomic classification and biodiversity assessment (Struck et al. 2018; Olivares et al. 2024; Peng et al. 2025). At the same time, hybridization can generate admixed individuals whose phenotypes overlap extensively with those of parental species, making reliable identification based on morphology alone difficult or impossible (Antoniou et al. 2018). Together, these processes pose major challenges for accurately delineating species boundaries and detecting introgression in natural populations(Yuan et al. 2025).

One such system occurs in marine environments, where the European lobster (*Homarus gammarus*) is increasingly threatened by hybridization with the invasive American lobster (*H. americanus*) following human-mediated introductions (Jørstad et al. 2011; Davies and Wootton 2018; Barrett et al. 2020). The introduced *H. americanus*, which is typically larger and more aggressive than the native *H. gammarus*, poses multiple risks to European lobster populations, including direct competition, predation, and the potential transmission of novel pathogens(Jørstad et al. 2011). In addition, interspecific hybridization raises concerns about maladaptive introgression and the long-term genetic integrity of native populations (Todesco et al. 2016).

Morphological traits commonly used to identify *H. americanus* or putative hybrids in European waters are not consistently diagnostic, particularly for admixed individuals and backcrosses (Jørstad et al. 2007; Jørstad et al. 2011)(Stebbing et al. 2012)(Jørstad et al. 2007; Jørstad et al. 2011). As a result, genetic tools have become central to monitoring the spread of non-native individuals and assessing the extent of hybridization. Jenkins et al. (Jenkins, Ellis, and Stevens 2019) developed a panel of 79 single nucleotide polymorphisms (SNPs), which was subsequently applied by Ellis et al. (Ellis et al. 2020) to reliably distinguish *H. gammarus, H. americanus*, and first-generation (F1) hybrids. However, the extent to which this panel can resolve introgression over longer timescales, such as first- and second-generation backcrosses, has not been systematically evaluated.

Here, we present an empirical resampling–based evaluation of a 79-SNP panel developed for hybrid detection in *Homarus* lobsters (Jenkins, Ellis, Triantafyllidis, et al. 2019; Ellis et al. 2020). Using genotype resampling from empirical parental populations to generate individuals with known ancestry proportions, we assess the panel’s resolution in distinguishing not only pure species and F1 hybrids, but also first- and second-generation backcrosses. Rather than relying on forward-time demographic simulations, our approach constructs *in silico* crosses directly from observed multilocus genotypes, thereby retaining the empirical allele frequency distributions and multilocus structure present in the original dataset, as commonly employed in validation studies of hybrid detection methods(Anderson and Thompson 2002; Vähä and Primmer 2006; Fitzpatrick 2012). While the European lobster (*H. gammarus*) and American lobster (*H. americanus*) serve as the biological case study, our framework and results speak more broadly to the challenges and possibilities of molecular introgression monitoring in applied conservation contexts.

## Materials and Methods

### Empirical SNP dataset

All analyses were based on an empirical SNP dataset comprising 1,591 individuals genotyped at 79 SNP loci, including *Homarus gammarus, H. americanus*, and putative hybrids (Jenkins, Ellis, Triantafyllidis, et al. 2019; Ellis et al. 2020). Genotypes were handled in R (v4.3.3) using the adegenet framework for genind objects (Jombart and Ahmed 2011). Individuals/populations corresponding to *H. americanus* (“Americanus” and “AmerCook”) were treated as the American parental reference. Atlantic European *H. gammarus* (all non-Mediterranean *H. gammarus* populations, excluding putative hybrids) were treated as the European parental reference. Mediterranean populations (Adr, Ale, Chi, Csa, Ion, Laz, Sar, Sky, Spo, The, Tor) and the empirical hybrid group (“HybridX”) were excluded from the parental pools used to generate resampled classes. No new sampling was conducted for this study, and no ethical approval was required.

### Empirical resampling framework for generating hybrid classes

To evaluate whether the 79-SNP panel can distinguish pure individuals and hybrids of known ancestry proportions, we generated synthetic genotypes by resampling from the empirical parental pools rather than forward simulation. This approach reconstructs expected genotype distributions under Mendelian segregation using observed allele-count data at each locus.

Genotypes were represented as allele-count vectors in the genind allele-column space. For each locus, a haploid gamete was generated from a sampled diploid individual by drawing one allele according to the allele counts at that locus (i.e., sampling an allele with probability proportional to its observed allele count in that individual at that locus). Two gametes were then combined to create a diploid offspring genotype by summing the allele indicators across loci.

Using this procedure, we generated seven “true” ancestry classes corresponding to expected proportions of *H. americanus* ancestry:

- 0: European parental class (resampled European individuals).
- 1: American parental class (resampled American individuals).
- 0.5 (F1): offspring of one European gamete and one American gamete.
- 0.25, 0.125: backcrosses toward Europe (F1 × Europe; then BC1 × Europe).
- 0.75, 0.875: backcrosses toward America (F1 × America; then BC1 × America).

For each class, we generated balanced datasets with equal sample size per class, using a grid of N per class = 20, 50, 100, 200, 500. For each N, we ran 100 independent replicates (each replicate using a unique random seed drawn from 1–100), regenerating all seven classes in each replicate. The resampled class datasets were combined into a single genind object for downstream clustering/assignment.

### Genetic assignment with snapclust including explicit hybrid classes

Genetic assignment was performed using snapclust (adegenet) with k = 2 parental clusters and hybrid modelling enabled (Beugin et al. 2018; Jombart and Ahmed 2011). Because snapclust labels the two parental clusters as “A” and “B” without intrinsic biological meaning, we treated A/B labels as returned by the model and evaluated accuracy against the corresponding expected hybrid class labels produced by snapclust under the specified hybrid coefficients.

Hybrid modelling was parameterised using a set of hybrid coefficients representing expected ancestry proportions. We used: hybrid.coef={0.125,0.25,0.5,0.75,0.875} constructed by taking the base set {0.125, 0.25, 0.5} and explicitly including complementary values (1 − x), reflecting first- and second-generation backcross expectations under simple Mendelian inheritance ensuring symmetric evaluation of backcrosses toward either parental cluster.

For each replicate dataset, snapclust returned posterior membership probabilities for the parental and hybrid classes. For evaluation, each individual was assigned to a single predicted class by selecting the class with maximum posterior probability.

### PCA and projection of resampled genotypes

We performed principal component analysis (PCA) on the empirical genotype matrix (allele-count columns from the genind@tab representation) after mean-imputation of missing values and standardization (centering and scaling). The PCA model (loadings/rotation, centering, and scaling parameters) was learned only from the empirical dataset. Resampled individuals of known ancestry were then projected into this fixed empirical PCA space by applying the same centering and scaling parameters and multiplying by the empirical PC loadings. As a result, empirical and resampled individuals share identical principal component axes, and differences in position reflect only their relative placement within the genetic variation captured by the 79-SNP panel.

### Statistical analysis

Assignment performance was quantified using two metrics:

Strict accuracy: an individual was counted as correctly assigned if its predicted class label exactly matched the expected snapclust label for its known (“true”) resampled class. Expected labels were defined for each true class (0, 0.125, 0.25, 0.5, 0.75, 0.875, 1) based on snapclust’s naming convention of hybrid classes.

Nearest-class accuracy: because adjacent hybrid classes can be difficult to separate with limited SNP numbers, we also computed a relaxed metric where an assignment was counted as correct if the predicted class belonged to an allowed set of the true class and its immediate neighbours along the ancestry gradient (e.g., 0.25 was considered correct if assigned to 0.125-, 0.25-, or 0.5-class labels; 0 was considered correct if assigned to 0 or 0.125; similarly for the American-side backcrosses). For each replicate and each *N*, we computed (i) overall accuracy across all individuals and (ii) per-class accuracy for each of the seven true classes. Summary statistics were obtained as the mean and standard deviation of replicate accuracies at each *N*.

All computations were performed in R (v4.3.3). Core dependencies for the resampling and snapclust pipeline included adegenet, dplyr, tidyr, and tibble. The analysis scripts and associated resources required to reproduce the pipeline are available in the project repository, together with the empirical data required to run the resampling-based evaluation.

### Use of Large Language Model

We used ChatGPT (version 5.2, OpenAI) to assist with code refactoring, language editing, and proofreading. All scientific interpretations, analyses, and conclusions are the responsibility of the authors.

## Results

### Classification accuracy across hybrid classes and sampling depth

For all hybrid classes, increasing the number of individuals sampled per class led to improvements in strict accuracy, with the most pronounced gains observed between 20 and 50 individuals per class. However, gains plateaued rapidly thereafter, with limited improvement beyond approximately 100–200 individuals per class. This pattern was observed for both European- and American-biased backcrosses, with reduced discrimination among adjacent hybrid generations persisting across sampling depths (**Figure 1**).

**Figure 1.**
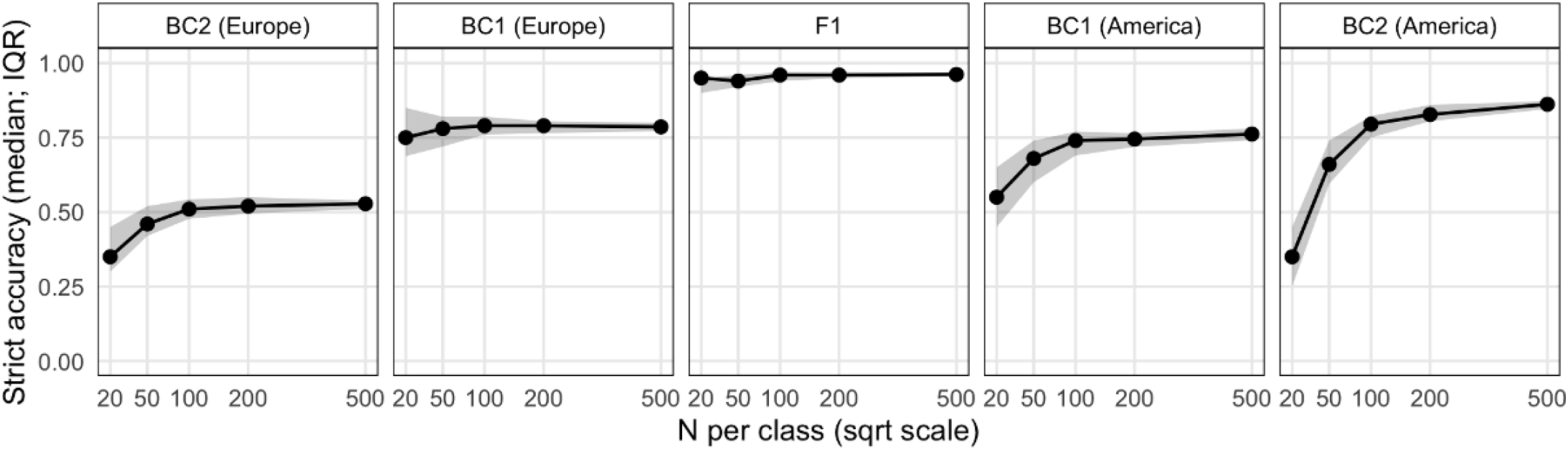
Strict classification accuracy across sampling depth for hybrid classes. Strict classification accuracy is shown as a function of the number of individuals sampled per class (*N* per class) across replicate resampling runs. Panels are faceted by the true resampled hybrid class: second-generation backcross to Europe (BC2 Europe), first-generation backcross to Europe (BC1 Europe), first-generation hybrid (F1), first-generation backcross to America (BC1 America), and second-generation backcross to America (BC2 America). Points and connecting lines indicate the median strict accuracy across replicates. Shaded ribbons represent the interquartile range (IQR), defined by the 25th and 75th percentiles (Q25–Q75) of accuracy across replicates. The x-axis is shown on a square-root scale.

Second-generation backcrosses exhibited the lowest strict accuracies across sampling depths, whereas first-generation backcrosses showed intermediate performance. By contrast, nearest accuracy remained consistently high (>0.99) across all classes and sample sizes. Most incorrect assignments involved neighbouring hybrid classes rather than distant ancestry categories.

### Projection of resampled hybrid genotypes onto empirical genetic space

Principal component analysis (PCA) was first performed on the empirical genotype dataset to characterize the major axes of genetic variation among lobster groups (**Figure 2**). The first two principal components explained 7.6% and 5.4% of the total genetic variance, respectively. Atlantic European and American individuals formed well-separated clusters primarily along PC1, whereas Mediterranean individuals were differentiated mainly along PC2. Empirical hybrids (HybridX) occupied intermediate positions between the Atlantic European and American clusters, consistent with admixed ancestry (Ellis et al. 2020).

**Figure 2.**
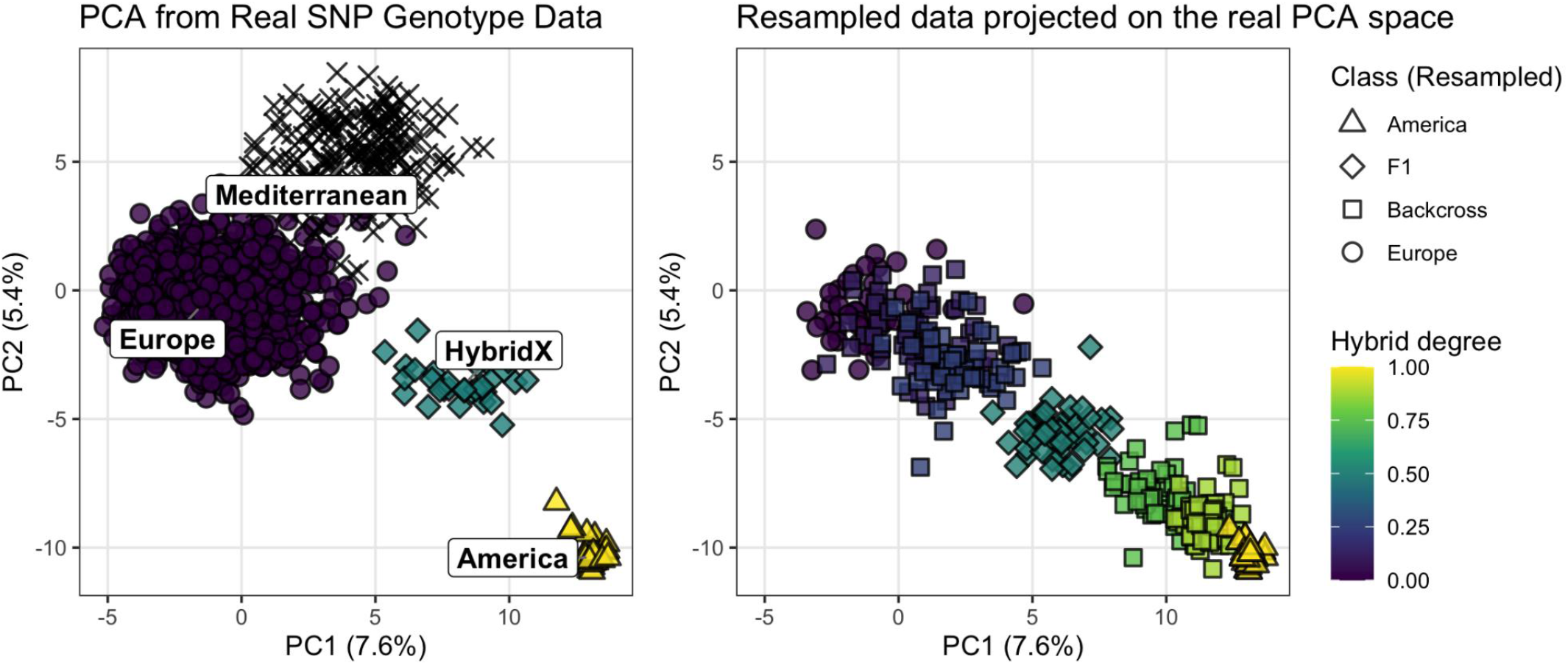
PCA of empirical and resampled lobster genotypes in a shared ordination space. The left panel shows the PCA result based on the empirical SNP dataset (79 loci), illustrating geographic origins (Europe, Mediterranean, HybridX, America). The right panel shows a projection of resampled individuals (a dataset with **N = 50 individuals per class** (seed = 1)) onto the empirical PCA space, and both panels share identical PC axes and variance explained (PC1 = 7.6%, PC2 = 5.4%). In the right panel, color gradient indicates the proportion of *H. americanus* ancestry (dark to light; 0 = European parental, 1 = American parental), with the sam color mapping applied to both left and right panels, while point shape in the projected panel denotes the true resampled class (Europe, backcross, F1 and America); Mediterranean individuals occur only in the empirical dataset for reference.

Resampled genotypes of known ancestry were then projected onto this fixed empirical PCA space, using the same centering, scaling, and loading parameters derived from the empirical data. Both empirical and resampled individuals are therefore displayed within the same coordinate system. Under this projection, resampled individuals formed a continuous gradient between the two parental clusters (**Figure 2**). Individuals with ancestry proportions close to 0 or 1 overlapped closely with the corresponding Atlantic European and American empirical clusters, whereas intermediate hybrid classes (ancestry 0.25, 0.5 and 0.75) occupied positions between these parental extremes.

**The ordering of resampled individuals along PC1 followed their ancestry proportions**, indicating that the primary axis of empirical genetic differentiation largely reflects the ancestry gradient between the two parental lineages (**Figure 2**). At the same time, intermediate hybrid classes showed substantial overlap in PCA space, particularly among neighbouring ancestry categories. This overlap corresponds to the distribution of snapclust assignments, where incorrect classifications occurred primarily between adjacent hybrid classes. Resampled hybrids overlapped with empirical HybridX individuals in PCA space, indicating that the resampled genotypes occupy regions of genetic space observed in the empirical dataset.

### Misclassification structure among hybrid classes

To examine the structure of classification errors in more detail, we visualised snapclust assignments for a representative resampled dataset with N = 50 individuals per class (seed = 1) as a confusion matrix (**Figure 3**). For each true resampled class, proportions of individuals assigned to each inferred class were calculated, such that rows sum to one.

**Figure 3.**
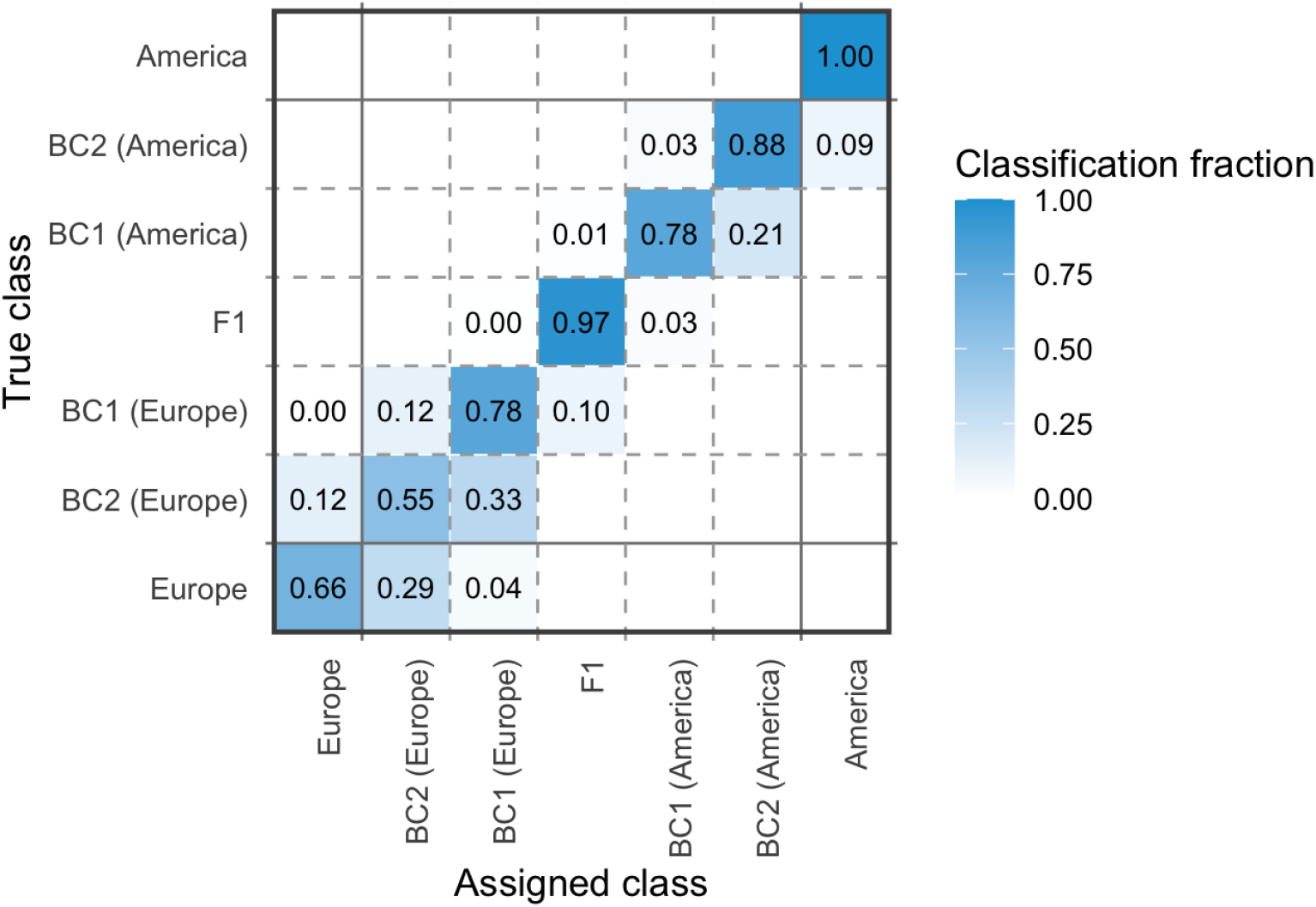
Confusion matrix of snapclust assignments across resampled hybrid classes. Snapclust assignment results for a representative resampled dataset with *N* = 50 individuals per class (seed = 1), visualised as a confusion matrix. The heatmap shows the within-class fraction of individuals assigned to each inferred class (x-axis) for each true resampled class (y-axis), with rows summing to one. Solid lines delineate the boundaries between pure parental classes (Europe and America) and the hybrid continuum, while dashed lines indicate transitions among hybrid classes.

Correct assignments were concentrated along the diagonal of the matrix, particularly for the parental classes (ancestry 0 and 1), which showed high assignment fidelity. In contrast, intermediate hybrid classes exhibited substantial off-diagonal signal, reflecting misclassification primarily among neighbouring ancestry categories. For example, individuals with true ancestry proportions of 0.25 and 0.125 were frequently assigned to each other or to the adjacent 0.5 class, while assignments to distant classes were rare.

Misclassifications occurred in both directions along the ancestry gradient, with no consistent asymmetry between European- and American-biased classes. Extreme backcross classes (0.125 and 0.875) showed partial overlap with their respective parental classes, consistent with the gradual blending of genotypes along the hybrid continuum. Overall, the confusion matrix indicates that reduced strict accuracy reflects ambiguity among adjacent hybrid classes rather than overlap between parental lineages.

### Calibration of posterior probabilities from snapclust

To assess whether posterior probabilities produced by snapclust reflect assignment reliability, we examined the relationship between maximum posterior probability and strict classification accuracy using a representative resampled dataset (**Figure 4**). Individuals were grouped into bins according to their maximum posterior probability, and strict accuracy was calculated within each bin.

**Figure 4.**
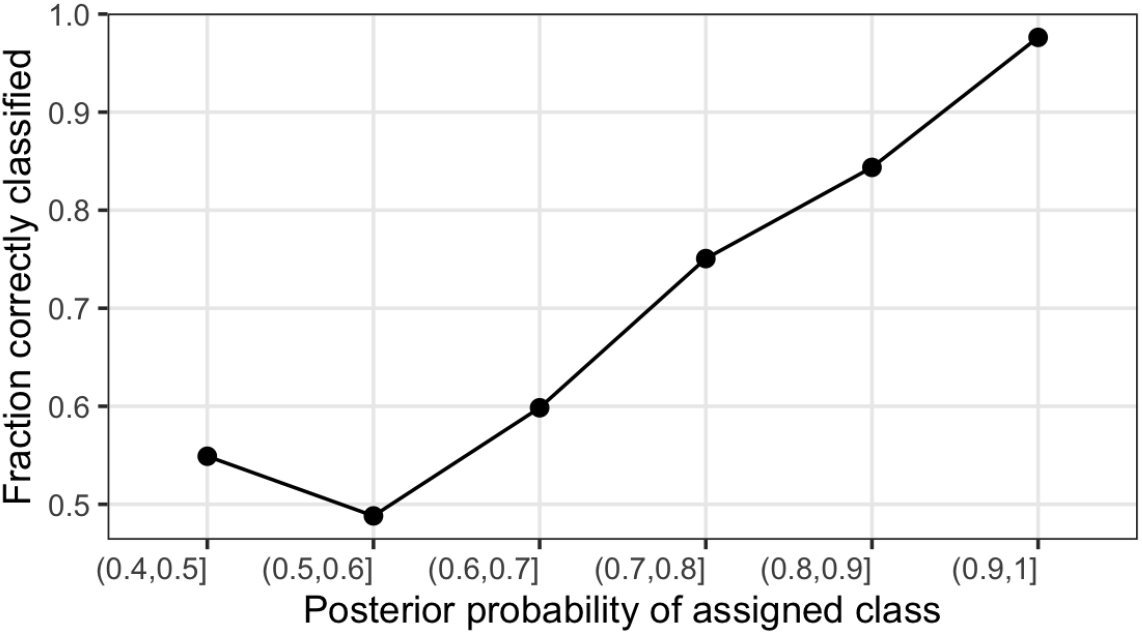
Relationship between snapclust posterior probabilities and classification accuracy. Strict assignment accuracy is shown as a function of the maximum posterior probability of the assigned class, grouped into probability intervals. Posterior probability bins are denoted using interval notation, where intervals of the form *(a, b]* include values greater than *a* and less than or equal to *b*. For each posterior probability bin, accuracy was calculated as the fraction of individuals whose assigned class exactly matched the true resampled class.

Strict accuracy increased monotonically with posterior probability. Assignments with posterior probabilities above 0.9 were associated with near-perfect strict accuracy, whereas assignments with lower posterior support showed substantially reduced accuracy. Individuals with posterior probabilities below 0.6 exhibited strict accuracies near 0.5, indicating considerable uncertainty in exact class assignment.

## Discussion

### Species boundaries, introgression, and the challenge of hybrid detection

Hybridization and introgression have long been central topics in evolutionary biology because they reveal that species boundaries often allow differential gene flow across parts of the genome rather than acting as absolute barriers (Harrison and Larson 2014). The concept of *semipermeable species boundaries*, whereby certain genomic regions remain differentiated in the face of hybridization while others are exchanged, has been repeatedly emphasized in hybrid zone research, particularly with the advent of molecular markers that enable fine-scale characterization of divergence and gene flow (Frade et al. 2010; Valencia-Montoya et al. 2020; Satokangas et al. 2023). This “genic view” highlights why morphological criteria alone often fail to capture underlying genetic structure, especially in cases of cryptic divergence and backcrossing. Accordingly, molecular tools that quantify ancestry gradients are necessary to delineate species identity and levels of introgression in natural populations(Massatti and Winkler 2022).

### Empirical performance of the 79-SNP panel and implications for ancestry resolution

Our results confirm that the 79-SNP panel robustly captures parental differentiation between *H. gammarus* and *H. americanus*, even at relatively low sampling depths per class. However, classification accuracy declines for intermediate hybrid categories, especially under strict assignment criteria, and improves with increased sample size.The decline in strict accuracy for intermediate classes reflects a general limitation of reduced SNP panels combined with discrete hybrid class models(Vähä and Primmer 2006): as the signature of introgressed ancestry becomes subtle, adjacent hybrid classes occupy overlapping regions of genetic space and are inherently difficult to distinguish (Baack and Rieseberg 2007). Consequently, while the 79-SNP panel reliably identifies broad ancestry classes, strict separation among multiple adjacent hybrid generations is less certain with limited numbers of loci; notably, subsequent extension of the SNP panel did not substantially alter the observed clinal pattern of genetic variation, as reported by Ellis et al. (2023), who found that the original marker set already captured the major axes of differentiation in this system .

### Integrating clustering, ordination, and probabilistic inference in hybrid analysis

The patterns of uncertainty observed in strict classification were mirrored by projections of resampled genotypes onto the principal component space defined by the empirical dataset: resampled individuals formed a smooth gradient between parental clusters, with substantial overlap among neighbouring hybrid categories. This concordance between ordination and assignment analyses suggests that the uncertainty among adjacent classes is rooted in the continuous nature of admixture at these marker densities.Importantly, posterior probabilities from snapclust were strongly associated with strict classification accuracy: high posterior support corresponded to near-perfect strict assignment, while lower posterior probabilities indicated greater uncertainty. Such probabilistic metrics therefore provide useful indicators of assignment confidence and can inform reporting thresholds in applied contexts. This approach aligns with broader practice in population genetics, where posterior support is increasingly used to quantify uncertainty in ancestry estimation (Oliveira et al. 2015; Chakraborty and Rannala 2023; Zbinden et al. 2023).

### Broader relevance, methodological context, and future directions

Reduced SNP panels have been applied in a variety of taxa beyond crustaceans. For example, panels of ancestry-informative markers have been developed to estimate introgression levels in honey bee subspecies, demonstrating that a relatively small set of markers can accurately assign individuals to lineage categories and estimate admixture proportions in conservation settings (Muñoz et al. 2015). Such studies highlight the practical value of reduced marker sets for routine screening in non-model systems where whole-genome resources are limited.Nevertheless, the challenges documented here,where finer discrimination among adjacent hybrid classes is sensitive to marker number and sample depth,are consistent with broader findings in hybrid zone research. Within the larger methodological landscape of hybrid detection, approaches range from ordination and clustering to genomic clines and whole-genome scans, each with distinct trade-offs between resolution and data requirements(Berger and Yu 2023). Empirical benchmarking, such as the resampling-based evaluation presented here, provides a realistic assessment of tool performance under conditions that approximate natural genotype distributions.

Finally, although this study does not directly assess adaptive consequences of introgression, work in diverse taxa, including plants, invertebrates and humans, has shown that introgressed alleles can sometimes confer adaptive traits across species boundaries (Hsieh et al. 2019; Valencia-Montoya et al. 2020; Termignoni-Garcia et al. 2022; Horta et al. 2025). Future research that integrates ancestry estimation with functional genomic data could further enrich understanding of hybrid dynamics and their evolutionary implications.

## Data Availability Statement

This study used both empirical and resampling-based genotype data.

All resampled datasets generated for the performance evaluation, together with all analysis scripts required to reproduce the results, are available in the project repository at: https://github.com/mariesaitou/paper_2025-/tree/main/lobster_resampling.

Empirical SNP genotype data were originally published by Ellis et al. (2020) and are included in the project repository with permission for non-commercial research use. No new raw sequence data were generated in this study.

## Acknowledgments

TWe thank fiskeridirektoratet for their support throughout the NMBU long-term lobster monitoring project. We thank Trond O. Haugen, Stein R. Moe, Jonathan E. Coleman, Odd Arne Sørensen, Knut Asbjørn Solhaug, and Linda Eikaas for their contributions to the NMBU long-term lobster monitoring program. We thank Lars Grønvold for an initial methodological suggestion regarding empirical resampling.

## References

Anderson EC, Thompson EA. 2002. A model-based method for identifying species hybrids using multilocus genetic data. Genetics. 160(3):1217–1229.

Antoniou A, Frantzis A, Alexiadou P, Paschou N, Poulakakis N. 2018. Evidence of introgressive hybridization between Stenella coeruleoalba and Delphinus delphis in the Greek Seas. Mol Phylogenet Evol. 129:325–337.

Baack EJ, Rieseberg LH. 2007. A genomic view of introgression and hybrid speciation. Curr Opin Genet Dev. 17(6):513–518.

Barrett CJ, Cook A, Stone D, Evans C, Murphy D, Johnson P, Thain M, Wyn G, Grey M, Edwards H, et al. 2020. A review of American lobster (Homarus americanus) records around the British Isles: 2012 to 2018. Hydrobiologia. 847(15):3247–3255.

Berger B, Yu YW. 2023. Navigating bottlenecks and trade-offs in genomic data analysis. Nat Rev Genet. 24(4):235–250.

Chakraborty S, Rannala B. 2023. An efficient exact algorithm for identifying hybrids using population genomic sequences. Genetics. 223(4). doi:10.1093/genetics/iyad011. http://dx.doi.org/10.1093/genetics/iyad011.

Chambers EA, Lara-Tufiño JD, Campillo-García G, Cisneros-Bernal AY, Dudek DJ Jr, León-Règagnon V, Townsend JH, Flores-Villela O, Hillis DM. 2025. Distinguishing species boundaries from geographic variation. Proc Natl Acad Sci U S A. 122(19):e2423688122.

Chan KO, Alexander AM, Grismer LL, Su Y-C, Grismer JL, Quah ESH, Brown RM. 2017. Species delimitation with gene flow: A methodological comparison and population genomics approach to elucidate cryptic species boundaries in Malaysian Torrent Frogs. Mol Ecol. 26(20):5435–5450.

Christie K, Strauss SY. 2019. Reproductive isolation and the maintenance of species boundaries in two serpentine endemic Jewelflowers. Evolution. 73(7):1375–1391.

Davies CE, Wootton EC. 2018. Current and emerging diseases of the European lobster (Homarus gammarus): a review. Bull Mar Sci. doi:10.5343/bms.2017.1142. http://dx.doi.org/10.5343/bms.2017.1142.

Ellis CD, Jenkins TL, Svanberg L, Eriksson SP, Stevens JR. 2020. Crossing the pond: genetic assignment detects lobster hybridisation. Sci Rep. 10(1):7781.

Ellis CD, MacLeod KL, Jenkins TL, Rato LD, Jézéquel Y, Pavičić M, Díaz D, Stevens JR. 2023. Shared and distinct patterns of genetic structure in two sympatric large decapods. J Biogeogr. 50(7):1271– 1284.

Fitzpatrick BM. 2012. Estimating ancestry and heterozygosity of hybrids using molecular markers. BMC Evol Biol. 12:131.

Frade PR, Reyes-Nivia MC, Faria J, Kaandorp JA, Luttikhuizen PC, Bak RPM. 2010. Semi-permeable species boundaries in the coral genus Madracis: introgression in a brooding coral system. Mol Phylogenet Evol. 57(3):1072–1090.

Harrison RG, Larson EL. 2014. Hybridization, introgression, and the nature of species boundaries. J Hered. 105 Suppl 1(S1):795–809.

Horta P, Raposeira H, Juste J, Razgour O, Rebelo H. 2025. Adaptive introgression as an evolutionary force: A meta-analysis of knowledge trends. Evol Appl. 18(6):e70103.

Hsieh P, Vollger MR, Dang V, Porubsky D, Baker C, Cantsilieris S, Hoekzema K, Lewis AP, Munson KM, Sorensen M, et al. 2019. Adaptive archaic introgression of copy number variants and the discovery of previously unknown human genes. Science. 366(6463). doi:10.1126/science.aax2083. http://dx.doi.org/10.1126/science.aax2083.

Jenkins TL, Ellis CD, Stevens JR. 2019. SNP discovery in European lobster (Homarus gammarus) using RAD sequencing. Conserv Genet Resour. 11(3):253–257.

Jenkins TL, Ellis CD, Triantafyllidis A, Stevens JR. 2019. Single nucleotide polymorphisms reveal a genetic cline across the north-east Atlantic and enable powerful population assignment in the European lobster. Evol Appl. 12(10):1881–1899.

Jombart T, Ahmed I. 2011. adegenet 1.3-1: new tools for the analysis of genome-wide SNP data. Bioinformatics. 27(21):3070–3071.

Jørstad KE, Agnalt A-L, Farestveit E. 2011. The Introduced American Lobster, Homarus americanus in Scandinavian Waters. In: In the Wrong Place - Alien Marine Crustaceans: Distribution, Biology and Impacts. Dordrecht: Springer Netherlands. p. 625–638.

Jørstad KE, Prodohl PA, Agnalt A-L, Hughes M, Farestveit E, Ferguson AF. 2007. Comparison of genetic and morphological methods to detect the presence of American lobsters, Homarus americanus H. Milne Edwards, 1837 (Astacidea: Nephropidae) in Norwegian waters. Hydrobiologia. 590(1):103– 114.

Massatti R, Winkler DE. 2022. Spatially explicit management of genetic diversity using ancestry probability surfaces. Methods Ecol Evol. 13(12):2668–2681.

Muñoz I, Henriques D, Johnston JS, Chávez-Galarza J, Kryger P, Pinto MA. 2015. Reduced SNP panels for genetic identification and introgression analysis in the dark honey bee (Apis mellifera mellifera). PLoS One. 10(4):e0124365.

Olivares I, Tusso S, José Sanín M, de La Harpe M, Loiseau O, Rolland J, Salamin N, Kessler M, Shimizu KK, Paris M. 2024. Hyper-Cryptic radiation of a tropical montane plant lineage. Mol Phylogenet Evol. 190(107954):107954.

Oliveira R, Randi E, Mattucci F, Kurushima JD, Lyons LA, Alves PC. 2015. Toward a genome-wide approach for detecting hybrids: informative SNPs to detect introgression between domestic cats and European wildcats (Felis silvestris). Heredity (Edinb). 115(3):195–205.

Peng J-C, He Z, Zhang Z-Q. 2025. Standing genetic variation and introgression shape the cryptic radiation of Aquilegia in the mountains of Southwest China. Commun Biol. 8(1):684.

Porto-Hannes I, Burlakova LE, Zanatta DT, Lasker HR. 2021. Boundaries and hybridization in a secondary contact zone between freshwater mussel species (Family:Unionidae). Heredity (Edinb). 126(6):955–973.

Sardell JM, Uy JAC. 2016. Hybridization following recent secondary contact results in asymmetric genotypic and phenotypic introgression between island species of Myzomela honeyeaters. Evolution. 70(2):257–269.

Satokangas I, Nouhaud P, Seifert B, Punttila P, Schultz R, Jones MM, Sirén J, Helanterä H, Kulmuni J. 2023. Semipermeable species boundaries create opportunities for gene flow and adaptive potential. Mol Ecol. 32(15):4329–4347.

Stebbing P, Johnson P, Delahunty A, Clark P, McCollin T, Hale C, Clark S. 2012. Reports of American lobsters, Homarus americanus (H. Milne Edwards, 1837), in British waters. Bioinvasions Rec. 1(1):17– 23.

Struck TH, Feder JL, Bendiksby M, Birkeland S, Cerca J, Gusarov VI, Kistenich S, Larsson K-H, Liow LH, Nowak MD, et al. 2018. Finding evolutionary processes hidden in cryptic species. Trends Ecol Evol. 33(3):153–163.

Termignoni-Garcia F, Kirchman JJ, Clark J, Edwards SV. 2022. Comparative population genomics of cryptic speciation and adaptive divergence in bicknell’s and Gray-cheeked thrushes (Aves: Catharus bicknelli and Catharus minimus). Genome Biol Evol. 14(1). doi:10.1093/gbe/evab255. http://dx.doi.org/10.1093/gbe/evab255.

Todesco M, Pascual MA, Owens GL, Ostevik KL, Moyers BT, Hübner S, Heredia SM, Hahn MA, Caseys C, Bock DG, et al. 2016. Hybridization and extinction. Evol Appl. 9(7):892–908.

Vähä J-P, Primmer CR. 2006. Efficiency of model-based Bayesian methods for detecting hybrid individuals under different hybridization scenarios and with different numbers of loci. Mol Ecol. 15(1):63–72.

Valencia-Montoya WA, Elfekih S, North HL, Meier JI, Warren IA, Tay WT, Gordon KHJ, Specht A, Paula-Moraes SV, Rane R, et al. 2020. Adaptive Introgression across Semipermeable Species Boundaries between Local Helicoverpa zea and Invasive Helicoverpa armigera Moths. Mol Biol Evol. 37(9):2568–2583.

Wu Y, Linan AG, Hoban S, Hipp AL, Ricklefs RE. 2024. Divergent ecological selection maintains species boundaries despite gene flow in a rare endemic tree, Quercus acerifolia (maple-leaf oak). J Hered. 115(5):575–587.

Yuan Y, Feng Y, Wang J, Ullah F, Yuan M, Gao Y. 2025. Integrative taxonomy for species delimitation: A case study in two widely accepted yet morphologically confounding Rosa species within sect. Pimpinellifoliae (Rosaceae). Mol Ecol. 34(21):e17779.

Zbinden ZD, Douglas MR, Chafin TK, Douglas ME. 2023. A community genomics approach to natural hybridization. Proc Biol Sci. 290(1999):20230768.

